# GenEditID: an open-access platform for the high-throughput identification of CRISPR edited cell clones

**DOI:** 10.1101/657650

**Authors:** Ying Xue, YC Loraine Tung, Rasmus Siersbaek, Anne Pajon, Chandra SR Chilamakuri, Ruben Alvarez-Fernandez, Richard Bowers, Jason Carroll, Matthew Eldridge, Alasdair Russell, Florian T. Merkle

## Abstract

CRISPR-Cas9-based gene editing is a powerful tool to reveal genotype-phenotype relationships, but identifying cell clones carrying desired edits remains challenging. To address this issue we developed GenEditID, a flexible, open-access platform for sample tracking, analysis and integration of multiplexed deep sequencing and proteomic data, and intuitive plate-based data visualisation to facilitate <u>gene</u> <u>edit</u>ed clone <u>id</u>entification. To demonstrate the scalability and sensitivity of this method, we identified KO clones in parallel from multiplexed targeting experiments, and optimised conditions for single base editing using homology directed repair. GenEditID enables non-specialist groups to expand their gene targeting efforts, facilitating the study of genetically complex human disease.

## BACKGROUND

In the last decade, there has been an explosion of data from the sequencing of human populations, including genome-wide association studies (GWAS) based on DNA microarrays and increasingly also whole exomes and whole genomes (1). These studies have revealed thousands of replicable genetic associations for complex diseases such as diabetes, obesity, Alzheimer’s Disease and breast cancer (2-4). However, mechanistically determining how these genetic associations contribute to disease remains challenging. Causal evidence requires careful functional follow-up experiments in model cellular systems, organisms and eventually in humans, but the traditional approach of characterising one gene at a time cannot keep pace with the rate of genetic discovery. Furthermore, many associated variants are non-coding, so the genetic elements responsible for conferring disease risk are often unclear (5, 6). This issue is exemplified by the fat mass and obesity associated (*FTO)* locus, in which intronic SNPs are strongly associated with obesity, largely due to increased food intake (7-10) irrespective of gender, age or ethnicity (11, 12). Despite intense study, the identify of the genetic elements that mediate SNP-associated phenotypes remains controversial. Some studies suggest that effect on appetite might not be driven by the *FTO* gene itself as initially thought, but instead by the nearby genes retinitis pigmentosa GTPase regulator-interacting protein-1 like (*RPGRIP1L*) (13-15), or by iroquois homeobox 3 (*IRX3*) and iroquois homeobox 5 (*IRX5*) (16, 17). Two powerful tools have recently emerged to help meet the challenge of uncovering disease mechanisms from the translating the growing wealth of genetic data: human pluripotent stem cells (hPSCs) and the CRISPR-Cas9 system (18, 19).

hPSCs facilitate human disease modelling since they can be indefinitely maintained in a pluripotent state and can theoretically be differentiated into any cell type in the body, including disease-relevant cell populations(20, 21). For example, hPSCs cell may be useful in dissecting which genes near *FTO* contribute to increased food intake since they can be differentiated into hypothalamic neurons that are pivotally important regulators of food intake and that express these candidate genes (22, 23). The CRISPR-Cas9 system enables most regions of the human genome to be efficiently edited. It consists of a ribonucleoprotein (RNP) complex, including a Cas9 nuclease that is targeted to specific regions of DNA by an approximately 20-base sequence within a guide RNA (gRNA) by forming a DNA-RNA hybrid with complementary DNA sequences (24-26). For Cas9 isolated from the bacterium *Streptococcus Pyogenes*, if the targeted DNA sequence contains a 3’ protospacer adjacent motif (PAM) of NGG, Cas9 will cleave the targeted DNA 3 bases 5’ to the start of the PAM site (27, 28) to create a double-strand break (DSB). The abundance of these PAM motifs in the genome allows most genes to be targeted by CRISPR-Cas9 (29). DSBs can be repaired by either the error-prone non-homologous end-joining (NHEJ) pathway which introduces either frame-preserving or frameshift mutations (30), or by the homology-directed repair (HDR) mechanisms which can be harnessed to introduce specific DNA alterations (31).

A major challenge in the field is how to effectively identify cell clones that have acquired desired edits. Next-generation sequencing (NGS) of multiplexed pools of amplicons provides an attractive solution to this problem, and CRISPR sequence analysis programmes built around this idea provide visualisations of mutation types (32-34) and frequency (35). However, to the best of our knowledge, there are no resources that provide a complete platform for amplicon generation and barcoding, sample tracking, sequence analysis, integration of distinct data forms (e.g. proteomic and sequencing), and intuitive visualisation to empower investigators in non-specialist labs to pursue high-throughput targeted gene editing.

To meet this challenge we developed a semi-automated, open-access, and user-friendly pipeline that captures the nature and frequency of CRISPR-Cas9-induced <u>gene</u> <u>edit</u>ing to <u>id</u>entify cell clones of interest, which we call GenEditID. Briefly, targeted regions of interest are amplified by PCR, barcoded, pooled and sequenced on an Illumina MiSeq. If genes of interest are expressed in the targeted cell type, protein expression data can be integrated with sequencing data to support the identification of knockout (KO) clones. Results are graphically represented to reflect the physical location of the clone on the plate, facilitating rapid and accurate clone recovery and further analysis. Using the *FTO* locus as an example, we provide a roadmap by which the community can use GenEditID to rapidly, affordably, and systematically explore genotype-phenotype relationships for genetically complex human diseases. Furthermore, we demonstrate how the sequencing depth and multiplexed nature of this approach enables targeted gene editing approaches to be optimised in cell populations before embarking on the laborious process of gene targeting and clone picking, allowing users to predict how many clones need to be picked in order to recover their clone of interest.

## RESULTS

We aimed to develop GenEditID to combine the strengths of laboratory information management system (LIMS)-based sample management with open-access customisable bioinformatic pipelines and a user-friendly graphical data display to facilitate the proliferation of parallel cellular gene editing experiments by non-specialist groups (Fig.1). GenEditID was implemented in Python, allowing basic experimental design and relevant sample details to readily be incorporated into a genome editing report. To establish this platform, we first turned to the estrogen receptor positive (ER+) breast cancer cell line MCF7, which is widely used to study breast cancer biology and is amenable to CRISPR-Cas9 gene editing (36). We targeted the oncogene signal transducer and activator of transcription 3 (*STAT3*), which is expressed in MCF7 cells but remains largely inactive in the absence of extracellular stimuli that trigger phosphorylation of tyrosine 705 (37). This approach allowed us to assess gene editing efficiency and identify successfully edited clones without confounding factors such as changes in the cell proliferation rate in response to deletion of *STAT3*, and to integrate mutually supportive data from protein expression and DNA sequencing.

**Figure 1).**
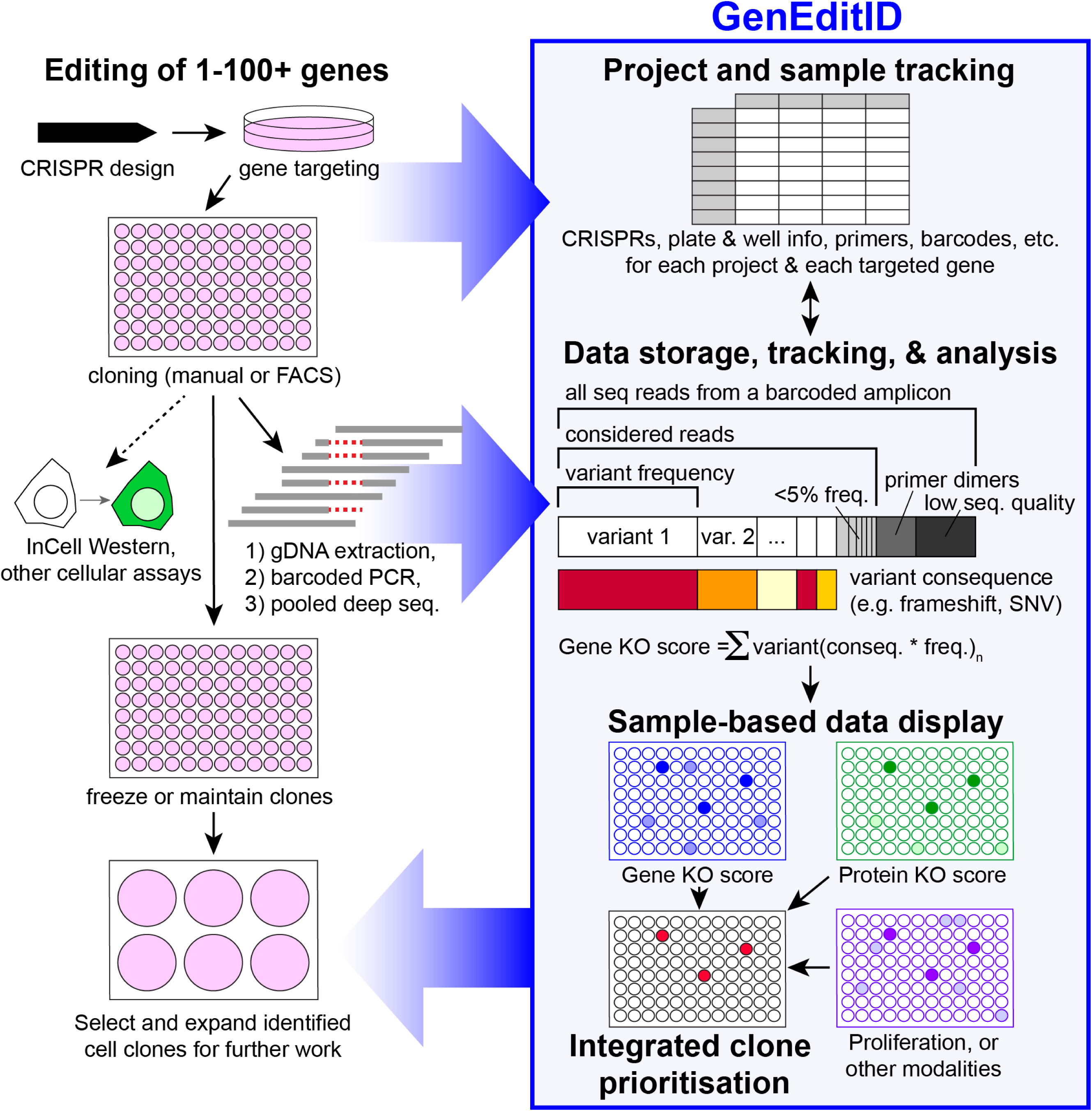
GenEditID facilitates the management and interpretation of multiplexed gene editing projects. After the design of CRISPR sequences to target one or multiple genes, sgRNAs are introduced along with Cas9 to a cell line of choice, which is then subjected to single-cell cloning by FACS or manual picking. The details of the CRISPR sequences, the location and plate that identifies each picked clone, the primers used to amplify the targeted region, any barcodes used to distinguish between PCR amplicons, and any other information pertaining to experimental conditions is entered into a centralised project and sample management system to track samples across projects. Next, data from assays across modalities (e.g. protein analysis by In-Cell Western and mutation analysis by barcoded PCR) feeds into GenEditID’s where it integrates with its sample tracking features and is analysed, for example to assess the burden of deleterious mutations carried by each clone (gene KOscore). Scores across these analysis modalities are graphically illustrated to facilitate the selection and expansion of cell clones for further analysis.

We designed four different gRNAs targeting exons 3 and 4 of *STAT3* (Fig. 2A, Supplementary Fig. S1A) that we cloned into a Cas9 expressing vector (pSpCas9(BB)-2A-GFP, PX458, Addgene#48138) (38), and separately transfected into a clonally-derived MCF7 cell line stably transfected with a vector expressing mStrawberry and luciferase (pCLIP-EF1a-LS). The use of a clonal cell line limited confounding factors associated with comparing clonal edited cell lines with a polyclonal parental cell line. Successfully transfected GFP+ single cells were distributed into 96 well plates using FACS, and clonal colonies were allowed to form. Viable colonies were consolidated into a new set of 96 well plates in duplicate, enabling one plate to be used to expand the clone for future use and the second plate to be used for clone characterisation. We first characterised 107 clones of cells by immunostaining wells with a *STAT3*-specific antibody, as well as a total cell stain used to normalise for differences in cell confluence in a high-throughput Li-Cor In-Cell Western (Fig. 2B) (39). Based on ratios of fluorescence intensity in channels corresponding to STAT3 abundance and total cell staining, we identified gene-edited clones with reduced STAT3 immunostaining relative to non-edited controls (Fig. 2B, white arrow). We validated loss of STAT3 protein for 8 such clones of interest using SDS-PAGE Western blotting (Supplementary Fig. S1B). Next, we tested whether the low protein expression observed in some targeted clones was due to CRISPR-Cas9-induced frameshifts in the *STAT3* genomic sequence and not spurious effects due to stress or clonal selection.

**Figure 2).**
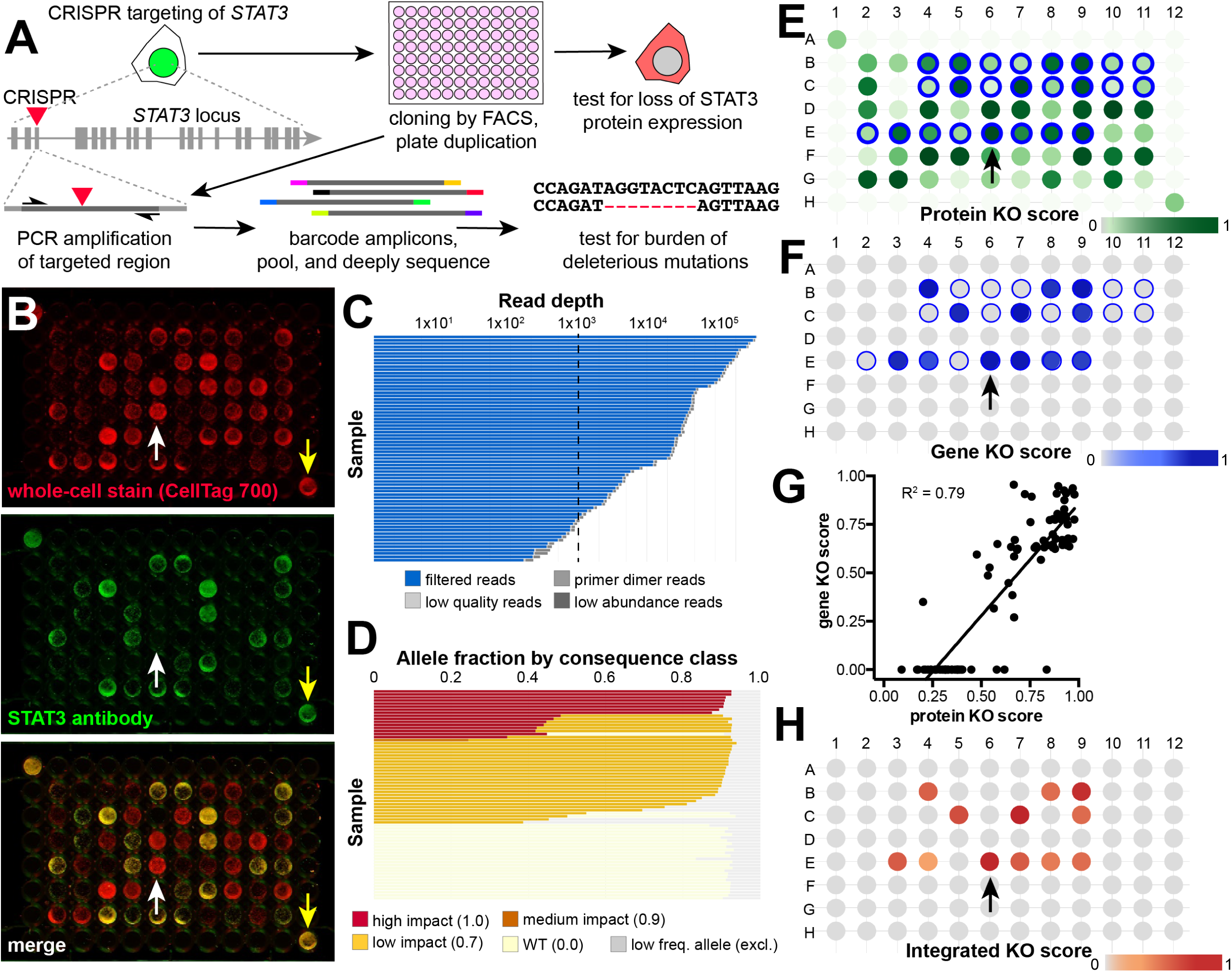
Integration of protein- and sequence-based information to inform knockout clone selection. **A)** Experimental schematic showing the CRISPR targeting of early constitutive exons of *STAT3* in the breast cancer cell line MCF7. After gene targeting, cells were clonally isolated by FACS into 96-well plates that were duplicated to facilitate clone propagation, protein analysis by In-Cell Western, and the barcoded PCR amplification of the targeted locus to facilitate deep sequencing and the identification of clones harbouring a high burden of deleterious (e.g. frameshift) mutations. **B)** Images from an In-Cell Western experiment where cells were stained for a whole-cell dye and with an anti-STAT3 antibody, and experiment clones were grown in the inner wells and wild-type controls in opposite corners (yellow arrows), revealing clones with reduced STAT3 expression (white arrow). **C)** Most samples have >1000 reads, providing good depth for assessing variant allele frequency. Low quality, low abundance, or primer dimer reads are discarded from subsequent analysis. **D)** Distribution of mutation frequencies and consequences across targeted cell clones used for calculating gene KO scores, where variants with an allele frequency of <5% (grey) do not contribute to these scores. **E**,**F)** Spatial heat maps of STAT3 protein KO score (E, compare to lower panel in B) and *STAT3* gene KO score. Note that only a subset of wells from E were analysed by Illumina sequencing (blue circles). **G)** Linear regression of STAT3 protein KO score and *STAT3* gene KO score shows a strong correlation (R^2^=0.79) between these independent measures of gene function. **H)** Heatmap of the integrated KO score (product of protein and gene KO scores).

To address this question, we extracted genomic DNA from 20 clones with low STAT3 protein expression and then PCR amplified and Sanger sequenced amplicons across the *STAT3* guide RNA target. We found that these clones indeed contained frameshift mutations disrupting both wild-type alleles of *STAT3* (Supplementary Fig. S1C). However, gene edited clones are often mosaic for a large number of alleles due to the persistence of CRISPR-Cas9 upon plasmid transfection (40, 41), and less abundant alleles are difficult to detect by Sanger sequencing. Therefore, we sought to test the extent to which sequencing information predicts STAT3 protein expression status across all clones by developing a bioinformatic pipeline for analysing sequencing reads for many clones in parallel. We reasoned that NGS would provide ample read depth and accuracy to permit multiplexing and the detection of low-abundance mutant alleles. We PCR amplified across the guide RNA target sites of *STAT3* for 96 clones, appended unique barcodes to each clone, pooled the barcoded amplicons, sequenced the pools, and bioinformatically identified amplicons arising from distinct cell clones (Supplementary Fig. S2).

Across these clones, we observed a median sequencing depth of 14,1322 reads corresponding to >90% of clones with at least 1000x coverage (Fig. 2C, Supplementary Table 3, Supplementary Fig. S3A), providing ample power to call mutation allele frequencies (Supplementary Fig. S3B). We next developed a sequence analysis pipeline to prioritize cell clones with a high burden of mutations predicted to result in gene loss of function (Fig. 1). To identify clones likely to have complete or near-complete gene KO, we aligned observed sequencing traces to the reference genome and quantified the number of reads corresponding to wild-type or variant sequence. We then omitted variants present at less than 5% abundance, classified remaining variant types (e.g. synonymous, missense, in-frame indels, frameshift indels) and assigned a score based on the likely consequence of each variant type on gene function (Fig. 2D, Supplementary Fig. S4, Supplementary Table 3). To determine the total burden of predicted gene-disrupting variants each clone, we calculated a “gene KO score” based on the aggregated product of mutant allele frequency and predicted mutation consequence (see Materials and Methods). To visualise both protein KO scores and gene KO scores, we implemented “heat maps” displaying these data based on the physical location on the plate for each clone (Fig. 2E and 2F). Note that due to differences in plate layout, only a subset of wells from plate 1 of this experiment (Fig. 2B) were submitted for Illumina sequencing (blue circles in Fig. 2E and F). Since some users may prefer to customise the calculation of KO scores or to integrate recently-developed methods for calling and classifying mutation types such as AmpliCan (35), we have made code developed for GenEditID freely available to the community at https://geneditid.github.io/

Next, we reasoned that the integration of gene KO and protein KO scores might provide stronger evidence to support KO clone selection. To test if mutations called by our sequence analysis pipeline predicts gene loss of function at the protein level, we compared protein KO and gene KO scores (Fig. 2G). We found that while control samples and clones identified as wild-type by sequence analysis tended to have similarly high STAT3 protein abundance, clones with a high burden of missense and frameshift mutations had significantly lower (R^2^ = 0.79, P < 0.0001) protein abundance, providing further confirmation of functional gene ablation. We therefore took the product of gene and protein KO scores to calculate an “integrated KO score” (Fig. 2H). These results indicate that count-based bioinformatic analysis of multiplexed NGS data predicts functional gene disruption. While multiple lines of evidence collected by high-throughput methods would be preferable to prioritise KO clone selection, this is often not possible since genes of interest may not be strongly expressed in the cell type used as the basis of gene editing, or appropriate antibodies may be lacking. For example, genes in the *FTO* locus that are implicated in obesity by GWAS are expressed in hypothalamic cells (42) but not are not highly expressed in hPSCs (23).

To test whether KO clones could be readily generated and identified across multiple genes in parallel in hPSCs, which are more challenging to edit than cancer cell lines (42, 43), we focused on the *FTO* locus. We first designed gRNAs to introduce double-strand breaks in early constitutive coding exons of the genes *FTO, RPGRIP1L, IRX3*, and *IRX5* (Fig. 3A) which are physically closest to the obesity-associated SNPs (Supplementary Fig. S5A). Next, we *in vitro*-transcribed four sgRNAs per gene, combined them with purified Cas9 protein, and tested their ability to cut PCR-amplified target DNA *in vitro* to select maximally active sgRNAs (Supplementary Fig. S5B and S5C). Since persistent Cas9 and sgRNA expression from plasmids can promote clone mosaicism and off-target activity (40, 41), and since double CRISPR-Cas9-induced strand breaks are cytotoxic to hPSCs (44, 45), we nucleofected hPSCs with Cas9 protein complexed with *in vitro* transcribed sgRNA, which has a half-life of approximately 24 hours in cells (46). hPSCs were re-plated at clonal density of 2 × 10^4^ cells/cm^2^ and when colonies emerged we manually picked them into one 96-well plate per targeted gene. After cells had reached approximately 70% confluence, we duplicated plates to allow one plate of clones to be frozen down for later use and the other to provide genomic DNA for our sequencing pipeline. This gene editing workflow is described in detail elsewhere (47).

**Figure 3).**
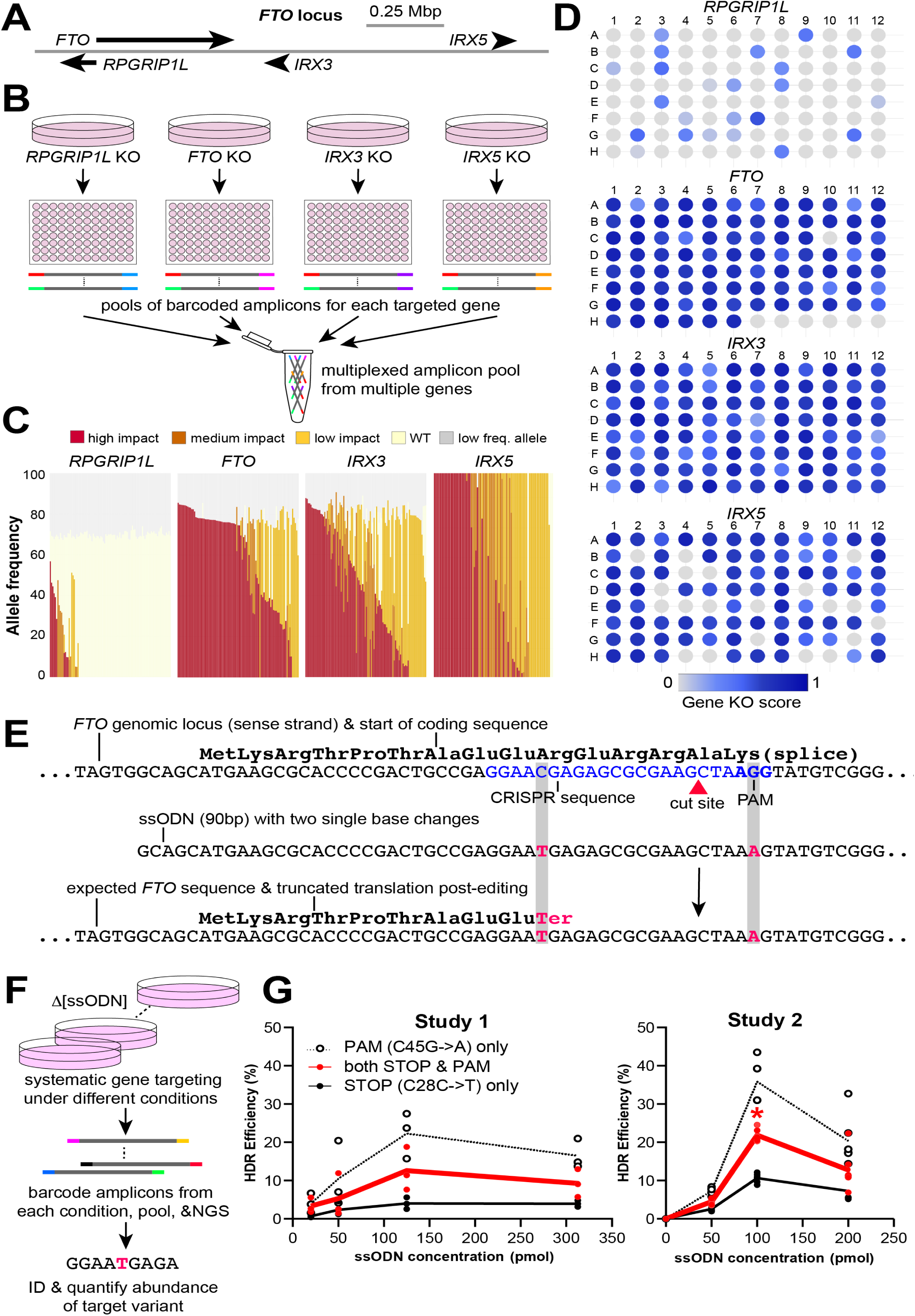
Multiplexed identification of knockout and knock-in clones by GenEditID. **A)** Schematic showing the spatial location of genes in the human *FTO* locus. **B)** Parallel KO of multiple genes in separate experiments, the generation of barcoded amplicons, and the pooling of these barcoded amplicons for multiplexed sequencing. **C)** Mutation allele frequency per targeted clone for each gene. **D)** Plate-based heat maps showing the location of clones with high gene KO scores (blue). **E)** Schematic of targeted ssODN-mediate mutation knock-in (red bases) in the first coding exon of *FTO*, to ablate the CRISPR PAM sequence (blue) and introduce and early stop codon (red) to ablate protein production. **F)** Schematic for the systematic modulation of CRISPR conditions and assessment of knock-in efficiency in un-cloned cell populations. **G)** Calculated knock-in efficiencies revealed a correlation with ssODN concentration that appeared to peak near 100-125 pmol. n=3 and n=6 for each concentration for Study 1 and Study 2, respectively.

Next, we PCR-amplified CRISPR target sites for each targeted gene and added distinct barcodes to each amplicon (Fig. 3B). The unique combination of amplicon and barcode allowed these 378 samples across 4 distinct amplicons to be combined into a “superpool” as previously described (48) along with an additional 987 samples from distinct experiments. We found that across these amplicons, there was no significant bias in sequencing coverage imposed by the barcodes (Supplementary Fig. S5D) and that the median sequencing coverage for these amplicons was 13,689 reads, with 96% of amplicons having 1000 or more reads (Supplementary Table 4). Using the pipeline we had built and tested using data from *STAT3* targeting, we categorised variant types and calculated gene KO scores to permit intuitive graphical data display (Fig. 3D), revealing efficient generation of KO clones for all targeted genes except *RPGRIP1L*. This web-based graphical output allows users to readily access data to retrieve clones of interest from highlighted wells for further analysis from remote locations, such as a tissue culture room.

In addition to NHEJ-mediated frameshifts, some groups may want to introduce specific mutations to test the consequence of disease-associated genetic variants or other functional elements. To complement NHEJ-mediated KO of *FTO*, we also introduced a single base mutation to introduce a premature stop codon into an early coding exon of *FTO* by HDR. To this end, we designed a single-stranded oligodeoxynucleotide (ssODN) with sequence complementarity to the target region but carrying the mutation of interest as well as a single-base mutation to ablate the PAM to eliminate further activity of CRISPR-Cas9 at the edited allele (Fig. 3E). Since HDR-mediated methods are inefficient, particularly in hPSCs (49), we harnessed the multiplexing capability and sequencing depth of NGS to optimise conditions for gene editing.

First, we identified optimal guides using *in vitro* cutting assays (Supplementary Fig. S5B). Next, we nucleofected hPSCs with CRISPR-Cas9 RNP and systematically increased concentrations of ssODN in biological triplicate. Rather than picking colonies, we extracted gDNA from the nucleofected hPSC populations, barcoded each treatment condition, and performed NGS (Fig. 3F). The sequencing depth provided by NGS allowed us to readily detect the desired edit, revealing that the fraction of correctly edited alleles varied with ssODN concentration, exceeding 10% allele frequency (approximately 20% of cells) for some conditions. However, the relationship between ssODN concentration and editing efficiency was non-linear and appeared to peak at a concentration around 100 pmol ssODN. This result was confirmed in a second independent experiment (Fig. 3G). We also tested the hypothesis that biotinylating Cas9 and adding a streptavidin tag to the biotinylating ssODN to physically link the CRISPR-Cas9 complex to the repair oligo would increase rates of HDR, as has been previously suggested (50). Despite clear Cas9 biotinylation, we found that this strategy unexpectedly abolished Cas9 activity (Supplementary Fig. S6A) and HDR (Supplementary Table 5), suggesting that future advances in gene editing could be rapidly tested and optimised using multiplexed NGS of targeted cell populations.

## DISCUSSION

We present GenEditID as a flexible, open-access pipeline that combines the benefits of a web-based project management system, a bioinformatic pipeline for data analysis, and a user-friendly graphical data output that allows efficient selections of clones of potential interest. Our aim was to reduce the expense, labour, and requisite expertise for groups (or core facilities) to carry out large-scale gene editing experiments. Here, we discuss the benefits of GenEditID, areas where it could be further developed, and observations about the nature of gene editing we observed in our dataset.

GenEditID can be readily customised and updated to incorporate new analysis tools (1) to meet diverse needs. We have therefore published the underlying code in the public domain https://geneditid.github.io/. In particular, the AmpliCount tool we developed was designed to rapidly analyse thousands of samples in parallel in order to prioritise clones for detailed follow-up analysis. Since sample information is clearly associated with raw sequencing traces by our sample tracking system, users can readily extract and further analyse data from clones of interest at the sequence level (51) or with more specialised bioinformatic tools (32-35), or incorporate these into GenEditID in lieu of AmpliCount.

Our analysis with AmpliCount revealed that most hPSC clones did not have simple distribution of heterozygous, compound heterozygous, or homozygous edited alleles but instead had a more complex mixture of alleles across a wide range of frequencies (Fig. 3C, Supplementary Table 4). These “mixed clones” were most likely generated by CRISPR-Cas9 activity that persisted across several cell divisions soon after transfection, as previously described by several groups (23, 40), despite the fact that we used a ribonucleoprotein complex of Cas9 and *in vitro*-transcribed sgRNA, which has a much shorter half-life in cells than plasmid- or virally-encoded Cas9 (46). These findings highlight the need to carefully analyse NGS data of candidate clones for the presence of mosaicism, and if necessary to perform a round of subcloning to isolate the desired clone.

Another advantage of the multiplexed barcoded amplicon sequencing pipeline of GenEditID is that the high sequencing depth enables scalable, multiplexed analysis. In this study, we ran 150 bp paired-end reads (300 cycles) on an Ilumina MiSeq. Assuming a minimum desired read depth of 1000x and one full 96 well plate of clones per gene, >10 or >100 genes could be screened in parallel using the MiSeq nano or standard v2 kits, respectively at a sequencing cost of less than 10 cents per clone. The establishment of a GenEditID pipeline at an institution would allow amplicons from different research groups to be pooled, facilitating cheaper and/or more frequent clone analysis. PCR barcoding by liquid handling robots could further increase the throughput and decrease the cost of this approach. The web-based data visualisation tool enables quick analysis of large numbers of clones and modularly integrates data such as protein expression, cell growth, or allelic frequency (Fig. 1), increasing confidence in clone selection.

The scalability of CRISPR-Cas9-based gene editing combined with streamlined clone selection provides a path to uncover the functional roles of disease-associated genes. This catalogue of genes is rapidly growing due to the proliferation and increased sample size of human population sequencing studies (52) In addition, the emergence of new methods for differentiating hPSCs into diverse cell types promises to enable the interrogation of gene function in disease-relevant cell populations. To illustrate this point, we targeted genes in the *FTO* locus, which was the first locus associated with obesity risk loci by GWAS (53, 54). The locus was subsequently linked to increased energy intake (7-10) but the identity of the genetic elements at this locus (or elsewhere) that mediate obesity risk remains controversial. Expression QTL studies of human cerebellum associate the obesity-linked SNPs to *IRX3* expression (17), but since the cerebellum is not an area of brain normally recognised to be involved in the control of food intake, there is a clear need to analyse brain regions pivotally important for body weight regulation such as the hypothalamus (55), which can now be generated from gene-edited hPCSs (22, 23). In this study, we report >80% mutation efficiency in target genes at the *FTO* locus in hPSCs, of which approximately 50% are frameshift mutations (Fig. 3C and 3D, Supplementary Table 4), demonstrating both the efficiency of the gene editing method and the sensitivity of GenEditID to detect these edited clones. For example, here we detected a clear dose-dependent effect of ssODN concentration on the efficiency of the targeted introduction of a point mutation in *FTO*. As CRISPR-Cas9 technology inexorably develops, we propose that new techniques can be more readily optimised using the tools described here to quantify low-frequency gene editing events within cell populations.

## CONCLUSION

We developed GenEditID to be a flexible, open-access platform to enable groups to track their samples, analyse and integrate amplicon sequencing and proteomic data. The combined data analysis is intuitively visualised to facilitate edited clone identification. The highly multiplexed approach enables the cost-effective and semi-automated identification of targeted clones and provides a powerful platform to systematically investigate complex human diseases in relevant cell types. We further show that GenEditID can be used to rapidly optimise conditions for editing single bases using homology directed repair. Using *FTO* as a proof of concept, we show how the platform can help the community to begin to elucidate the mechanisms by which genetic variants contribute to human disease.

## METHODS

### Cell lines and routine cell culture

The clonal MCF7 breast cancer cell line expressing mStrawberry and luciferase (a gift from Scott Lyons, Cold Spring Harbor Laboratory NY) was grown in DMEM (41966-029, Gibco) supplemented with 10% fetal bovine serum (FBS, 10500-064, Gibco), 50 U/ml penicillin and 50 ug/ml streptomycin (15070-063, Gibco) and 2mM L-glutamine (25030, Gibco) in a humidified 37°C incubator with 5% CO_2_. The HUES9 human embryonic stem cell line was grown on tissue culture plates coated wth Geltrex (Thermo Fisher Scientific) in mTeSR1 media (StemCell Technologies) and maintained in a humidified 37°C incubator with 5% CO_2_. Medium was changed every 24 hours. Cells were passaged with 1 mM ETDA for routine maintenance in mTeSR media supplemented with 10 μM ROCK inhibitor Y-27632 dihydrochloride (DNSK International). Please see below for culture details during gene editing. The absence of mycoplasma was confirmed using a EZ-PCR Mycoplasma Test Kit (Supplementary Fig. S6B ; Biological Industries, 20-700-20) following the manufacturer’s instructions.

### CRISPR-Cas9-mediated targeting of *STAT3*

Four different gRNAs with high predicted on-target and low predicted off-target activity targeting exons 3 and 4 of STAT3 (NM_139276) were designed using deskgen (www.deskgen.com). These guides were ordered from Sigma Aldrich and cloned into pSpCas9(BB)-2A-GFP (PX458, Addgene #48138). A stably transfected clonal MCF7 cell line expressing mStrawberry and luciferase (pCLIIP-EF1-LS) was transfected with these vectors. Successfully transfected GFP+ cells were purified by FACS into 6 well plates and allowed to recover for ∼1 week. Single mStrawberry+ cells (since the transfection with CRISPR-Cas9 vector was a transient transfection, the cells had lost GFP expression at this point) were then distributed into multiple 96 well plates by FACS. After ∼3 weeks, viable clonal colonies were consolidated on new 96 well plates in duplicate, so that one plate could be used for characterising clones and the other plate could be used to expand clones for later use. All sgRNAs and primer sequences are provided in Supplementary Table 1.

### In-Cell Western for STAT3

Five days after seeding, the test plate used for clone characterisation was fixed in 3.7% formaldehyde for 20 min at room temperature and subjected to In-cell western using a STAT3-specific antibody (9139, Cell Signalling). Briefly, cells were permeabilised by washing 5x in TBS + 0.1% Triton X-100 (Fisher Scientific, BP151-100) for 5 min at room temperature and then blocked for 1 h with TBS Odyssey blocking buffer (Li-Cor biosciences, 927-50000). Cells were then incubated with a STAT3 antibody (9139, Cell Signalling) in blocking buffer + 0.1% tween-20 (P1379, Sigma Aldrich) for 1-2 hours at room temperature or overnight at 4°C on a shaker. After washing 5x with TBS+0.1% Tween-20 for 5 min, cells were incubated with secondary antibody (Goat anti-mouse, 926-32210, Li-Cor) and a 1:500 dilution of CellTag 700 total cell stain (Li-Cor, 926-41090) for 45min at room temperature on a shaker (50 ul/well). Cells were then washed 4x with TBS+0.1% Tween-20 for 5 min followed by a final wash in TBS. Plates were then analysed using the Odyssey CLx Imaging System (Li-Cor) to obtain measurements for both total cell confluency and STAT3 expression. Signal intensity for STAT3 staining (green channel) was then divided by the signal intensity for the CellTag 700 stain (red channel) for each well to determine a STAT3 abundance ratio, and the resulting ratios were normalised to produce a “protein KO score”. The mean of negative control values, representing background staining, was subtracted from all ratios, and the resulting scores were divided by the mean of positive (non-edited) intensity scores, and the resulting values were subtracted from 1 so that WT lines would have scores near 0 and KO lines would have scores near 1.

### Production and testing of *in vitro*-transcribed gRNA

CRISPR guide RNAs were designed to target early coding gene regions in constitutive exons of genes in the *FTO* locus using Wellcome Trust Sanger Institute Genome Editing tool (http://www.sanger.ac.uk/htgt/wge/) and the Feng Zhang laboratory’s CRISPR design tool (http://crispr.mit.edu/) to maximise on-target and minimize off-target activity. For the production of gRNAs, a 120 nucleotide oligo (Integrated DNA Technologies Inc.) including the SP6 promoter, gRNA sequences, and scaffold region were used as a template for synthesis by *in vitro* transcription using the MEGAscript SP6 kit (Thermo Fisher, AM1330) as previously described [19]. The resulting sgRNAs were purified using the E.Z.N.A miRNA purification kit (Omega Bio-tek, R7034-01), eluted in RNase-free water, and stored at −80°C.Since gRNAs vay in their efficacy, we designed at least four gRNAs per gene of interest and tested their relative cutting efficiencies in *in vitro* cleavage assays as previously described (47). We selected the gRNAs that showed activity at the lowest Cas9 concentration at each target gene for transfection in the hPSC cells. All sgRNAs and primer sequences are provided in Supplementary Table 1.

### Cas9 protein production

Cas9 proteins were purified by the laboratory of Marko Hyvönen (University of Cambridge) from E. coli expressing *Streptococcus Pyogenes* Cas9 carrying a C-terminal fusion to a hexa-histidine tag from the pET-28b-Cas9-His plasmid (Addgene http://www.addgene.org/47327) (56). The soluble Cas9 protein was purified by a combination of nickel affinity and cation exchange chromatographies. The purified protein was concentrated to approximately 30 μM (4.8 mg/ml) in 20 mM HEPES pH 7.5, 500 mM KCl and 1% sucrose buffer and flash frozen for storage at −80°C.

### CRISPR-Cas9 ribonucleoprotein (RNP) complex-mediated editing in hESCs

For gene knockout by NHEJ, 3 μg purified sgRNA was mixed with 4 μg Cas9 protein (final volume <5 μl) for 10 min at room temperature to form stable RNP complexes. The complex was then transferred to a 20 μl single-cell suspension of 2 × 10^5^ hESCs in P3 nucleofection solution and electroporated using Amaxa 4D-Nucleofector™ (Lonza) with program CA137. Transfected cells were seeded onto Geltrex-coated 10 cm dishes containing a pre-warmed 1:1 mix of mTeSR1 and hESC medium containing 20% knockout serum replacement (KOSR) and 100 ng/ml bFGF, supplemented with 10 μM ROCK inhibitor. Rock Inhibitor was withdrawn after 24 hours. Single colonies were isolated manually 7-10 days after transfection and seeded into Geltrex-coated 96-well plates in 1:1 medium plus ROCK inhibitor, which was withdrawn after 24 hours. A total of 96 individual colonies were picked for each targeted gene, and maintained in 1:1 medium for 10-14 days. Once clones were close to confluent, each of the 96 well plates were duplicated by EDTA passaging to allow parallel cell cryopreservation and genomic DNA extraction as previously described (47). Briefly, cell cryopreservation was performed in hESCs culture media containing a final concentration of 40% FBS and 10% DMSO. Cells were slowly frozen in ice-cold freezing media using Mr. Frosty™ Freezing Container (Thermo Fisher Scientific, 5100-0001). During the thawing of hESCs, media was supplemented with CloneR (StemCell Technologies, 05889) for 72 hours.

### Generation and sequencing of pooled amplicons

Genomic DNA was extracted using HotShot buffer as previously described (47). The target regions were amplified from gDNA using locus-specific primers to generate amplicons approximately 150-200 bp in length as previously described (47). These “first-round” primers contained universal Fluidigm linker sequences at their 5’-end with the following sequences: Forward primer: 5’-acactgacgacatggttctaca -3’, Reverse primer: 5’-tacggtagcagagacttggtct-3’. Specifically, 20 μl PCR reactions were set up in 96 well plates using 1U FastStart high fidelity polymerase (Roche, 3553361001), 2 μl of extracted gDNA as template, 2 μl 10x HF buffer without MgCl_2_, 0.2 mM dNTPs, 0.2 μM primers, and 4.5 mM MgCl_2_, and run on the following programme: 95°C 2 min, followed by 36 cycles of (95°C 20 sec, 64.4°C 20 sec, 72°C 15 sec), 72°C 3 min. In the second round of PCR (indexing PCR), Fluidigm barcoding primers were attached to the amplicons to uniquely identify each clone. 2 μl linker PCR product diluted 1:10 was transferred to another 96-well PCR plate to perform this indexing PCR in 20 μl reactions containing 0.04 μM of Fluidigm barcoding primers (Supplementary Table 1), 2 μl 10x HF buffer without MgCl_2_, 0.2 mM dNTPs, 4.5 mM MgCl_2_, and 1U FastStart high fidelity polymerase (Roche, 3553361001). The PCR programme was 95°C 2 min, 16 cycles of (95°C 20 sec, 60°C 20 sec, 72°C 25 sec), 72°C 3 min. For sequencing library preparation, barcoded PCR products were combined in equal proportion based on estimation of band intensity on a 2% agarose gel, and the combined pool of PCR products was purified in a single tube using Ampure XP beads (Beckman-Coulter, A63880) at 1:1 (V/V) to the pooled sample, and eluted in 25 μl of water according to the manufacturer’s instructions. Library purity was confirmed by nanodrop, and final library concentration was measured using the Qbit fluorometer and diluted to 20 nM. Pooled libraries could be combined with other library pools adjusted to 20 nM, and the resulting “superpool” volume was adjusted to a final volume of 20 μl before sequencing.

### Introduction of STOP codon in FTO through HDR-mediated repair

To target a STOP codons to an early coding exon of FTO, we designed a ssODN template of 90 bp in length to be homologous to the target site but to contain single base mismatches to introduce a STOP codon and to ablate the PAM site to prevent re-cutting by CRIPSR/Cas9 as previously described (47)(Supplementary Table 1). ssODNs were synthesized (Integrated DNA Technologies Inc.), dried, and re-suspended in nuclease-free sterile water to a final concentration of 100 μM. Various amounts of ssODNs ranging from 20 pmol to 312.5 pmol were added to RNP complexes for nucleofection as described above. Editing efficiency was determined by sequencing the targeted locus using primers outside of the ssODN at the cell population level rather than in single picked colonies, and then counting the number of reads corresponding to the WT amplicon sequence, or sequences with one or both desired edits.

### Project tracking within the GenEditID web framework

We designed a Python-based web framework (web app) of GenEditID to facilitate the tracking of different projects, the tracking of samples within a project, and to facilitate the plate-based data integration and visualisation to help users identify clones of interest. When initiating a project, the user first creates a project via the web app (for a screenshot example of the web app home page see Supplementary Fig. S7) along with comments about the project purpose and design, and then submits an Excel configuration file containing the plate and well each sample originated from, the CRISPR sequences used, the primers and barcodes used for NGS analysis, and any other pertinent information (Fig. 1, Supplementary Table 2). This project information is later accessible via the web app and also programmatically, and enables samples to be uniquely identified so that data from different analysis modalities (e.g. sequencing, growth rate, protein abundance) can be loaded into the web app for tracking, integration, and visualisation. We designed the web app to enable integration with a laboratory information management system (LIMS) to automatically trigger a sequencing request when sample information is uploaded. Links to specific LIMS are intentionally omitted from the code published here since we anticipate that different institutions will wish to implement GenEditID within existing systems. Analysis can then be run outside the web app using setup scripts generated from the database, as described in further detail below.

### Amplicon analysis from NGS data with AmpliCount

The analysis of barcoded PCR amplicons is performed outside of the web app using modular scripts (https://geneditid.github.io/) that are adaptable to each user’s specific requirements. First, FASTQ files associated to the project are retrieved and configuration files are created to link sequencing information with the sample information stored in the GenEditID project configuration file. These de-multiplexed FASTQ files are then either merged, or joined using fastq-join if the target size is larger than the read length.

To analyse reads, we developed “ampli_count” (Supplementary Fig. S4), a tool that first finds amplicons using primer pairs and group variants with same sequence. Then we identified and filtered out reads of low quality across a 5bp sliding window (average read quality score < 10). Putative primer dimers were defined as any sequence smaller than the combined size of the forward and reverse primers, plus 10bp, and discarded. To focus downstream analysis on variants that reflect CRISPR-Cas9-induced edits rather than sequencing artefacts, sequences supported by 60 or fewer reads were also discarded. After obtaining “filtered reads” that passed these criteria, amplicon-specific variants were identified using the tool “variant_id” to determine variant type and consequence per site by pairwise alignment to human reference genome ensembl_grch38 using pairwise2 from Biopython (https://github.com/biopython/biopython), and using varcode (https://github.com/openvax/varcode) and pyensembl (https://github.com/openvax/pyensembl) to determine consequence. Since the aim of this analysis was to identify clones that carried a high burden of variants likely to lead to loss of gene function rather than generate a comprehensive description of variants observed upon gene editing, only variants with an overall frequency of 5% or higher were retained for downstream analysis. All filter steps and thresholds are tuneable by the user.

Remaining variants were classified according to their predicted consequence (Supplementary Fig. S4). Variants with multiple predicted consequences following sequence alignment were labelled “Complex”, or “ComplexFrameShift” if they contained a frameshift. Consequences were then given an impact weighted score based on their predicted effect on gene function, ranging from 0 for wild-type sequences to 1 for the gain of a premature stop codon. Variants were then grouped by similar consequence categories and these combined frequencies were multiplied by an impact weighting score and summed across all consequence categories to yield a “gene KO score” for each allele, where a score of 0 would correspond to all WT sequences, and a score of 1 would correspond to all predicted deleterious variants (Supplementary Fig. S4). Variant classification and weighting for KO score calculation can be readily altered via csv file.

### Clone score integration and data visualisation

After computing protein KO scores and gene KO scores, data were loaded back into the GenEditID database to facilitate their integration with stored sample information. Where both scores were available, the “integrated KO score” was calculated by taking the product of the gene and protein KO scores. To facilitate the selection and expansion of candidate KO clones, information about each clones’ plate and well position was used to graphically display computed scores as a “heat map” in 96 well plate format.

### Statistical analyses

Statistical analyses were performed using Graph Pad Prism version 8.1.0. A two-tailed Student’s *t*-test was performed to compare knock-in efficiencies among different conditions for variable amount of ssODNs. Unless otherwise stated, data shown represent the results of at least three independent experiments. P-values < 0.05 were considered significant.

## List of abbreviations

GWAS: genome-wide association studies
FTO: fat mass and obesity associated
RPGRIP1L: retinitis pigmentosa GTPase regulator-interacting protein-1 like
IRX3: iroquois homeobox 3
IRX5: iroquois homeobox 5
hPSCs: human pluripotent stem cells
RNP: ribonucleoprotein
gRNA: guide RNA
PAM: protospacer adjacent motif
DSB: double-strand break
NHEJ: non-homologous end-joining
HDR: homology-directed repair
NGS: Next-generation sequencing
KO: knockout
LIMS: laboratory information management system
ER_+_: estrogen receptor positive
STAT3: signal transducer and activator of transcription 3
ssODN: single-stranded oligodeoxynucleotide
KOSR: knockout serum replacement
Bio-Cas9: biotinylated form of Cas9

## DECLARATIONS

### Ethics approval and consent to participate

Not applicable

### Consent for publication

Not applicable

### Availability of data and material

The datasets generated and analysed during the current study have been deposited in NCBI’s Sequence Read Archive at https://dataview.ncbi.nlm.nih.gov/object/PRJNA543767?reviewer=ktlduo7ptjcsmrajrhnfnj0su2 (knockout of human *STAT3*);https://dataview.ncbi.nlm.nih.gov/object/PRJNA543845?reviewer=h1v7go700g7n1ocftv3hckrk3m (multiplexed knockout of human *RPGRIP1L, FTO, IRX3*, and *IRX5*); and htps://dataview.ncbi.nlm.nih.gov/object/PRJNA545266?reviewer=9ptviqm1j6bpfubtjdherr4r08 (targeted editing of human *FTO*).

### Authors’ contributions

YX, YCLT, and FTM conceived the project and wrote the manuscript with contributions from all other authors. RS generated data from *STAT3* targeting with the guidance of JC. YX and YCLT generated data from gene knockout and targeted gene editing of *FTO* and neighboring genes with the guidance of FTM. AP and CSRC generated the bioinformatics plots. AP, CSRC, AB and RAF developed bioinformatic tools under the guidance of ME and AR.

## Acknowledgements

We thank Magdalena Jura and Peter Kirwan for their assistance with culturing hESC lines, Beata Blaszscyk and Marko Hyvönen for generating and providing purified Cas9 and Cas9-biotin proteins, Kelly Holmes for generating MCF7 cells stably expressing luciferase and mStrawberry, and Sir Professor Stephen O’Rahilly for his generous financial support.

## SUPPLEMENTARY FIGURES

**Supplementary Figure S1. Validation of GenEditID with the clonal selection of *STAT3* KO lines. A)** A schematic diagram depicting the site-specific CRISPR designs to target exon 3 or 4 of the human *STAT3* gene. **B)** STAT3 protein expression by Fluorescent Western blot confirmed that STAT3 (green) was effectively knocked down in the MCF7 cells. Actin (red) was used as loading control. **C)** Comparison of growth rate with loss of STAT3 protein expression (ratio to total cell stain) and presence of insertion/deletion mutations (indels) in Sanger sequencing data. **D)** Sanger sequencing at the target site showing sequence chromatograms for clone B6 from plate 4 (right) showing an “A” insertion introduces a premature STOP codon. Clone E2 from plate 1 (left) as an example of STAT3 wild type at the corresponding locus.

**Supplementary Figure S2. PCR amplicon barcoding and pooling for NGS. A)** Genomic DNA is extracted from cell clones 96 well plates that were duplicated from another plate which is maintained or frozen. **B)** The CRISPR-targeted region is PCR amplified with locus-specific primers containing universal (L1 and L2) linker sequences (orange box). **C)** In a second PCR cycle (blue box), one of 96 unique Fluidigm forward barcodes are added to the end of the PCR product. To further increase the multiplexing capacity, up to 96 distinct reverse barcodes can be used. The resulting barcoded amplicons are then concentration normalised, pooled, and sequenced on the Illumina MiSeq in the presence of PhiX to increase library diversity.

**Supplementary Figure S3. Amplicon sequencing coverage and criteria used for variant quality control.** Log_10_-transformed distribution of read depth (A) and variant frequency (B) at one of the two PCR amplicons generated at the human *STAT3* locus, showing criteria for inclusion for downstream analysis: minimum read depth >1000 after filtering (A), and minimum variant allele frequency >5% (B).

**Supplementary Figure S4. Schematic flowchart of steps involved in variant calling and annotation by AmpliCount.** Throughout this flowchart, green indicates data outputs, orange indicates tools or file inputs, and processing steps are indicated in white. FASTQ files are first demultiplexed and reads are combined by merging or joining, depending on amplicon length (please see Materials and Methods). The “ampli_count” tool then filters out reads with low sequence quality, reads likely corresponding to primer dimers, and reads corresponding to low abundance (e.g. <60 reads) sequences to retain filtered reads from which variant frequencies are computed. The “variant_id” tool then removes all reads that have low abundance (e.g. <5%) relative to all filtered reads to streamline downstream alignment and variant classification steps. Remaining variants are pairwise-aligned, classified by variant consequence, scored as indicated. Variants in the same consequence categories are combined, and the combined frequencies are multiplied by the consequence scores and summed to yield gene KO scores for each clone (see also Fig. 1). The resulting data outputs include plots and tables, including graphical visualisation of gene KO scores in plate-based heat maps.

**Supplementary Figure S5. Multiplexed analysis of CRISPR targeting of the human *FTO* locus. A)** SNPs within intron 1 of *FTO* have a strong association with obesity, implicating the causal involvement of several nearby genes. **B)** Representative gel image of an *in vitro* cleavage assay evaluating gRNA:Cas9-mediated target cleavage. The arrowheads indicate nuclease cleaved products. **C)** Schema of the CRISPR RNA targeting early conserved coding exons in four candidate genes in the *FTO* locus. The CRISPR recognition sequence is shown in red and the PAM sequence is shown in blue. **D)** Log_10_-transformed sequencing depth per barcode of the four targeted sequenced and analysed in parallel indicating ample sequence quality and depth for variant analysis.

**Supplementary Figure S6. A)** Results from an *in vitro* Cas9 cutting assay where a biotinylated form of Cas9 (Bio-Cas9) was mixed with a biotinylated-ssODN and streptavidin was added at increasing concentrations to physically link Cas9 and the ssODN repair oligo, but this approach appeared to disrupt cutting ability, as indicated by the lower molecular weight bands observed with non-biotinylated (WT) Cas9. **B)** All hESC cell lines were tested for mycoplasma before and after gene editing and found to be negative.

**Supplementary Figure S7.** A screenshot of an example of the homepage of the Web App in the web browser to facilitate the tracking of different projects. Some key features of the Web App including “Create project” which enables the creation of a new project. “edit” allows the uploading of the configuration file that are subsequently loaded into the database to facilitate sample tracking. “view” displays target amplicon and a list of the samples. “ngs data” house all the resulting data outputs including the csv tables and plots.

## SUPPLEMENTARY TABLES

**Supplementary Table 1. Compilation of primer, barcode, CRISPR and ssODN sequences used in this study.** The table contains all the sequence information used in all the studies covered in this manuscript, including the CRISPR guide RNA sequences, target primer sequences used for the *in vitro* Cas9 cutting assay and target primer sequences for the first round (linker) PCR for NGS. The ssODN sequence is complementary to the *FTO* target region and carries mutations to introduce a premature STOP codon and a mutation to disrupt the PAM site to prevent further CRISPR-Cas9 activity.

**Supplementary Table 2. Example of a configuration file used to submit and track samples.** An example of a configuration file that are submitted by the user when initiating a project that contains all the relevant information from the user including the CRISPR sequences used, primers sequences for NGS analysis and sample coordination on the plate. All the information in the configuration file is later accessible via the web app allowing samples to be uniquely identified for tracking, integration and visualisation.

**Supplementary Table 3. Amplicon data from the human *STAT3* locus generated with AmpliCount.** “**Config STAT3**” contains the information including the target genomic location and the two sets of primers use to identify the two amplicons of interest. “**variantid STAT3**” contains the total reads after each step of the filtering process to remove reads with low quality, short length (primer dimers), or low abundance, and provides information on the variant frequency, type and consequence to give a variant score where the score is the consequence weight × variant frequency. “**impact STAT3**” tablet the combined frequency of all variants with identical consequence categories (high/medium/low impact). Finally, the “**koscores_amplicon**” provides individual table for each amplicon (i.e. STAT3 exon3 & exon4) on variants grouped in each consequence categories (i.e. variant with identical impact weighing) and their combined frequencies to yield a gene KO score where the score is the sum of impact weight × impact frequency for each allele and a score of 1 would correspond to all deleterious variants.

**Supplementary Table 4. Amplicon data from the human *FTO* locus generated with AmpliCount.** “**Config FTO plus**” contains the genomic location and the primer pairs use to identify the four target gene of interest, namely *FTO, IRX3, IRX5* and *RPGRIP1L.* “**variantid FTO plus**” table contains all total reads for each barcode and sequences reads after each step of the filtering process such as reads with low quality, primer dimers, low abundance, and lists the variant type and consequence to give a variant score. “**impact FTO plus**” shows data on the combined frequency of all variants with identical consequence weighing and categories the consequence into high/medium/low impact. “**koscores_amplicon**” tablet each of the four amplicons included in this study and provides data on variants grouped by same consequences categories and their combined frequencies to yield a gene KO score for each allele where a score of 1 would correspond to all deleterious variants.

**Supplementary Table 5. Targeted editing with ssODNs to introduce targeted mutations into *FTO*.** “**config ssODN**” lists the four types of variant sequence that were tested for, namely the wild-type *FTO* sequence, the sequence with both target sites mutated, the variant sequence with target site only where a STOP codon was introduced in the early coding exon of FTO and sequence where only the PAM site was mutated. “**amplicount Study 1**” shows the total and filtered (e.g. matching target sequences named above) sequencing reads in eight cell populations gene edited with either WT Cas9 or Cas9-biotin proteins (for ssODN concentration of 20 pmol, 50 pmol, 125 pmol and 312.5 pmol; n=3 for each condition). “**amplicount Study 2**” shows the total and filtered sequencing reads in another independent four groups of un-cloned cell populations (for ssODN concentration of 0 pmol, 50 pmol, 100 pmol and 200 pmol; n=6 for each condition). “**KoIN Study 1**” and “**KoIN study 2**” shows the calculated knock-in efficiencies for Study 1 and Study 2 respectively where the % of HDR efficiency were represented by the percentage of specific variant filtered read over the sum of all filtered reads for each barcode.

