## Supplementary figures and images for "GenEditID: an open-access platform for the high-throughput identification of CRISPR edited cell clones"

### Supplemental Figure1

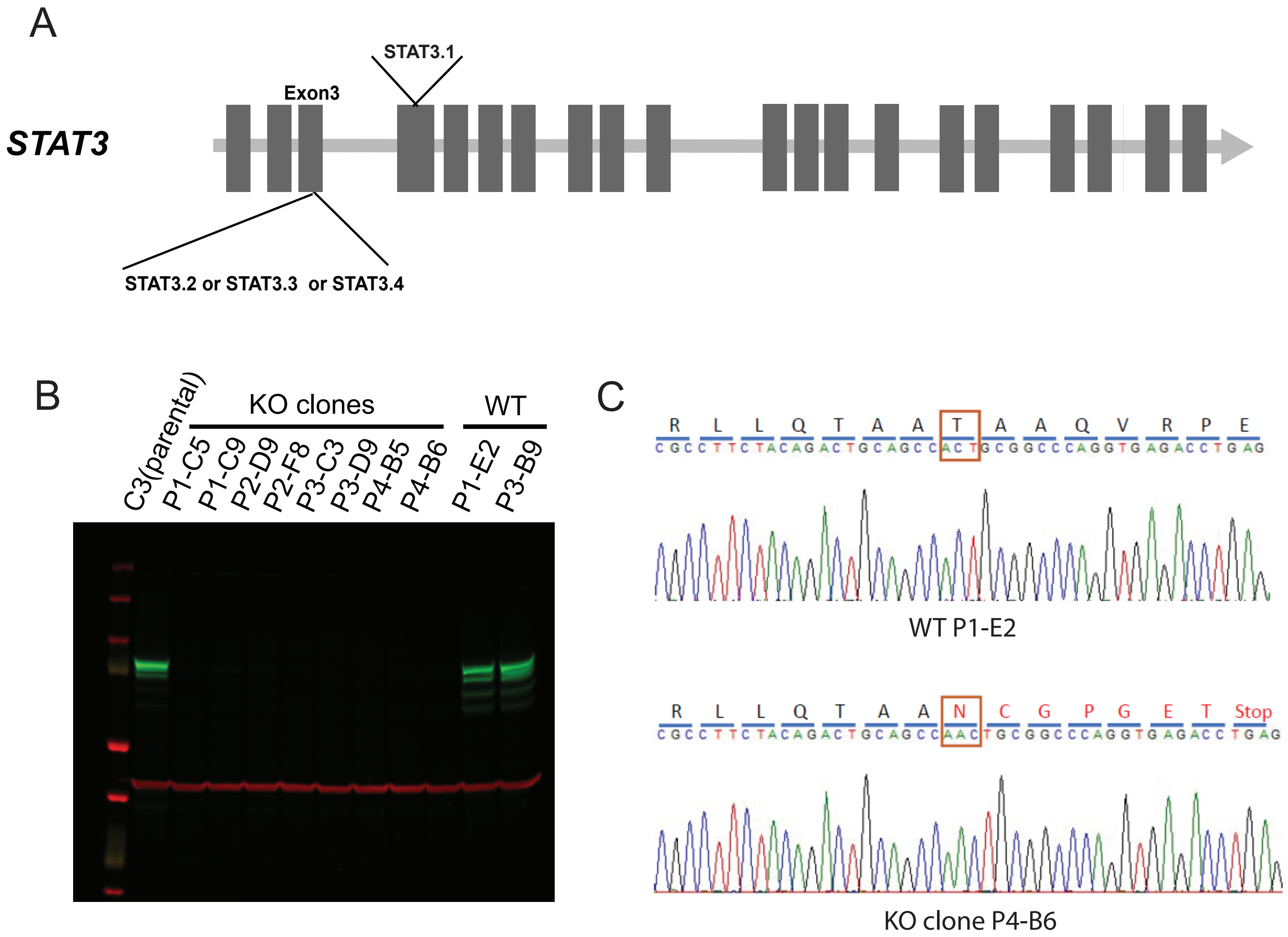


Supplementary Figure S1

### Supplemental Figure2

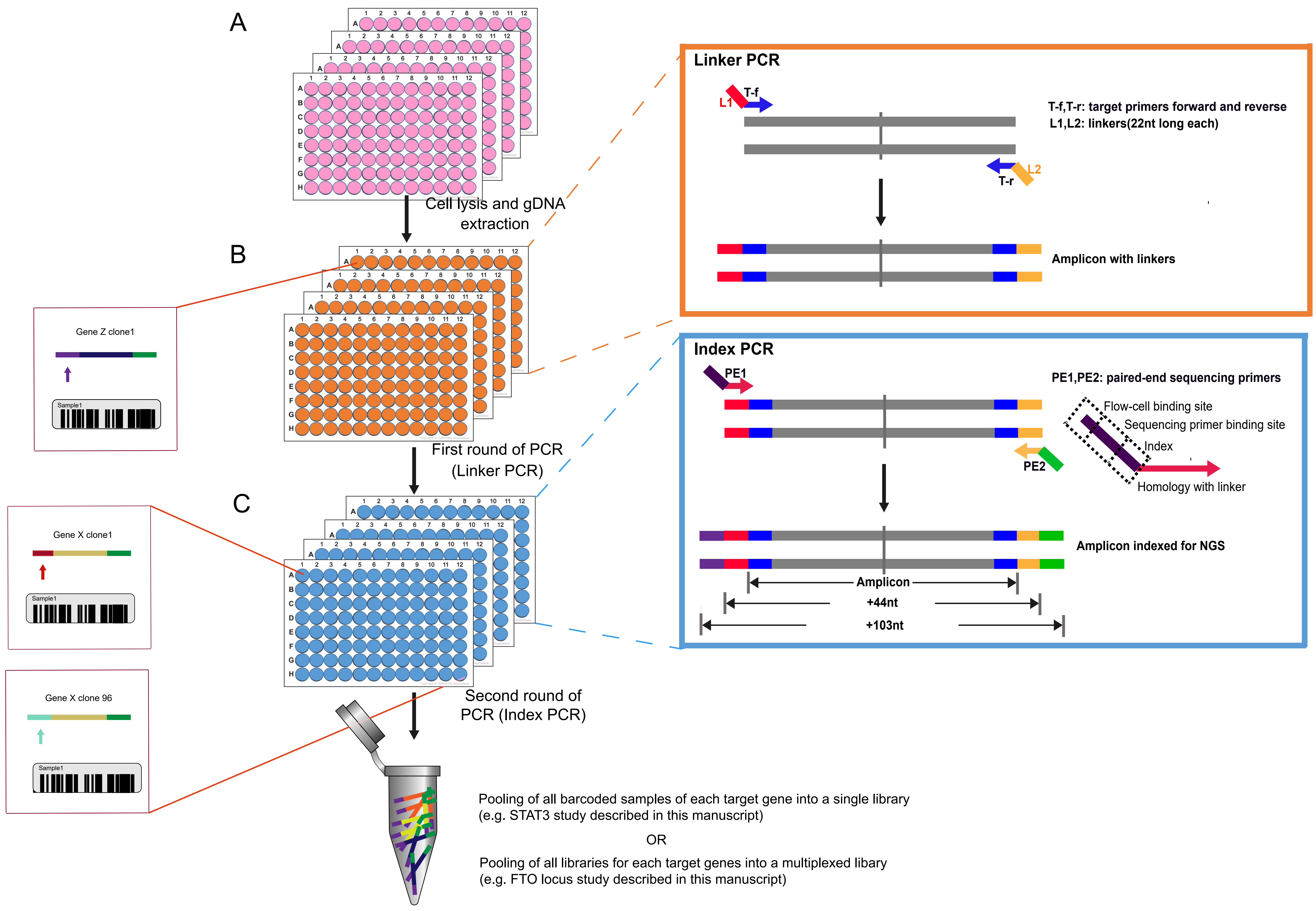


Supplementary Figure S2

### Supplemental Figure3

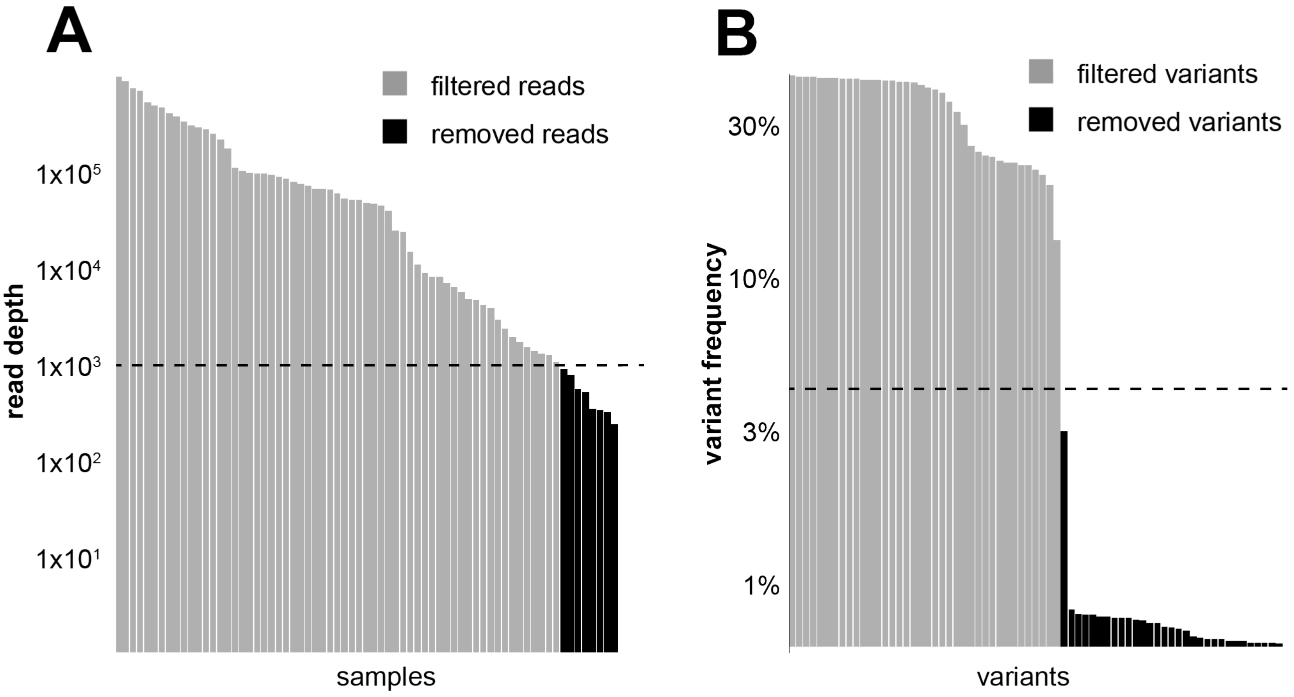


Supplementary Figure S3

### Supplemental Figure5

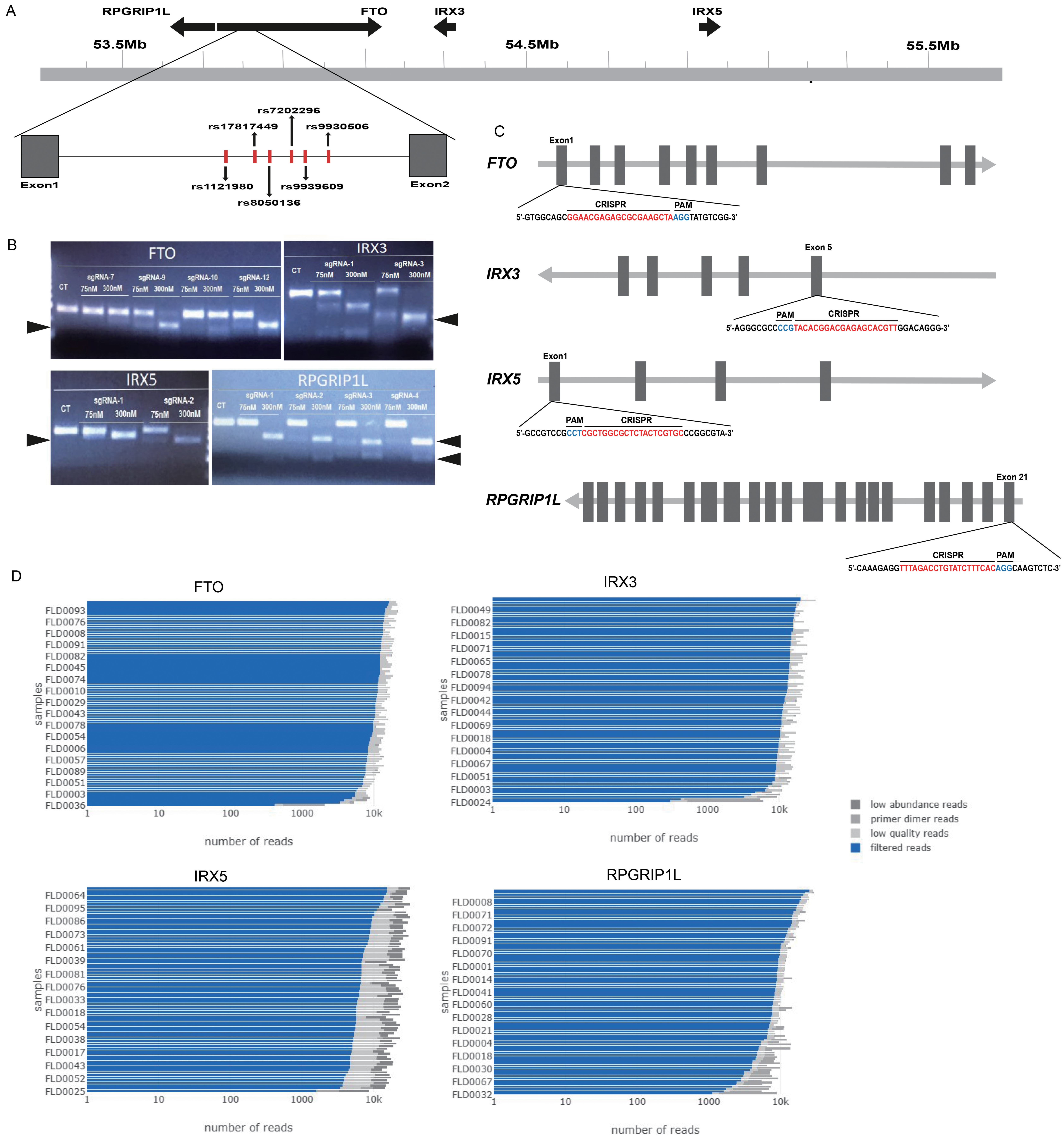


Supplementary Figure S5

### Supplemental Figure7

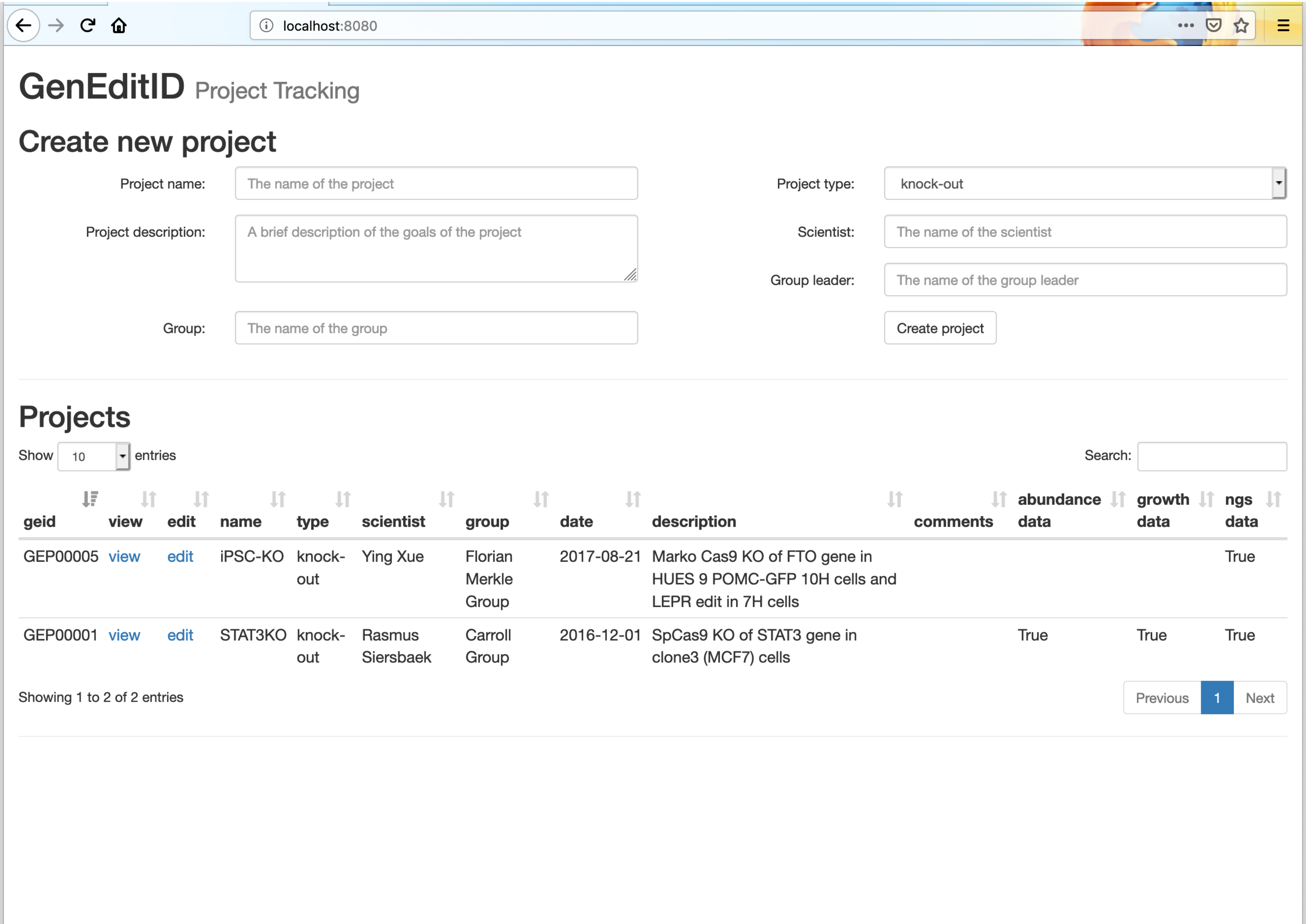


Supplementary Figure S7
