## Supplemental Figure4 for "GenEditID: an open-access platform for the high-throughput identification of CRISPR edited cell clones"

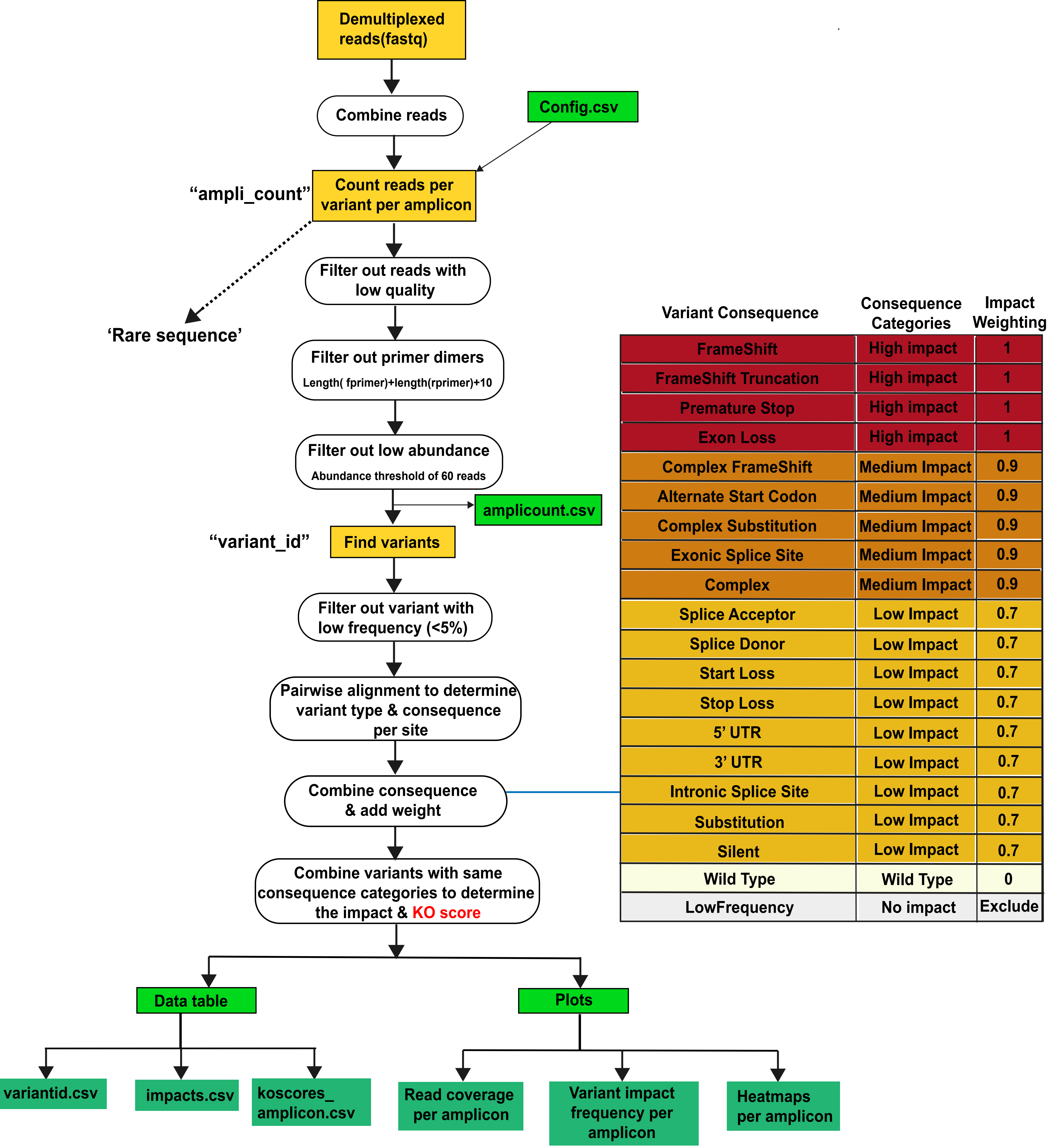


| Variant Consequence | Consequence Categories | Impact Weighting |
|---|---|---|
| FrameShift | High impact | 1 |
| FrameShift Truncation | High impact | 1 |
| Premature Stop | High impact | 1 |
| Exon Loss | High impact | 1 |
| Complex FrameShift | Medium Impact | 0.9 |
| Alternate Start Codon | Medium Impact | 0.9 |
| Complex Substitution | Medium Impact | 0.9 |
| Exonic Splice Site | Medium Impact | 0.9 |
| Complex | Medium Impact | 0.9 |
| Splice Acceptor | Low Impact | 0.7 |
| Splice Donor | Low Impact | 0.7 |
| Start Loss | Low Impact | 0.7 |
| Stop Loss | Low Impact | 0.7 |
| 5' UTR | Low Impact | 0.7 |
| 3' UTR | Low Impact | 0.7 |
| Intronic Splice Site | Low Impact | 0.7 |
| Substitution | Low Impact | 0.7 |
| Silent | Low Impact | 0.7 |
| Wild Type | Wild Type | 0 |
| LowFrequency | No impact | Exclude |

Supplementary Figure S4
