## Supplemental Figure6 for "GenEditID: an open-access platform for the high-throughput identification of CRISPR edited cell clones"

**A In vitro cutting with Bio-ssODN, Bio-Cas9 and streptavidin**

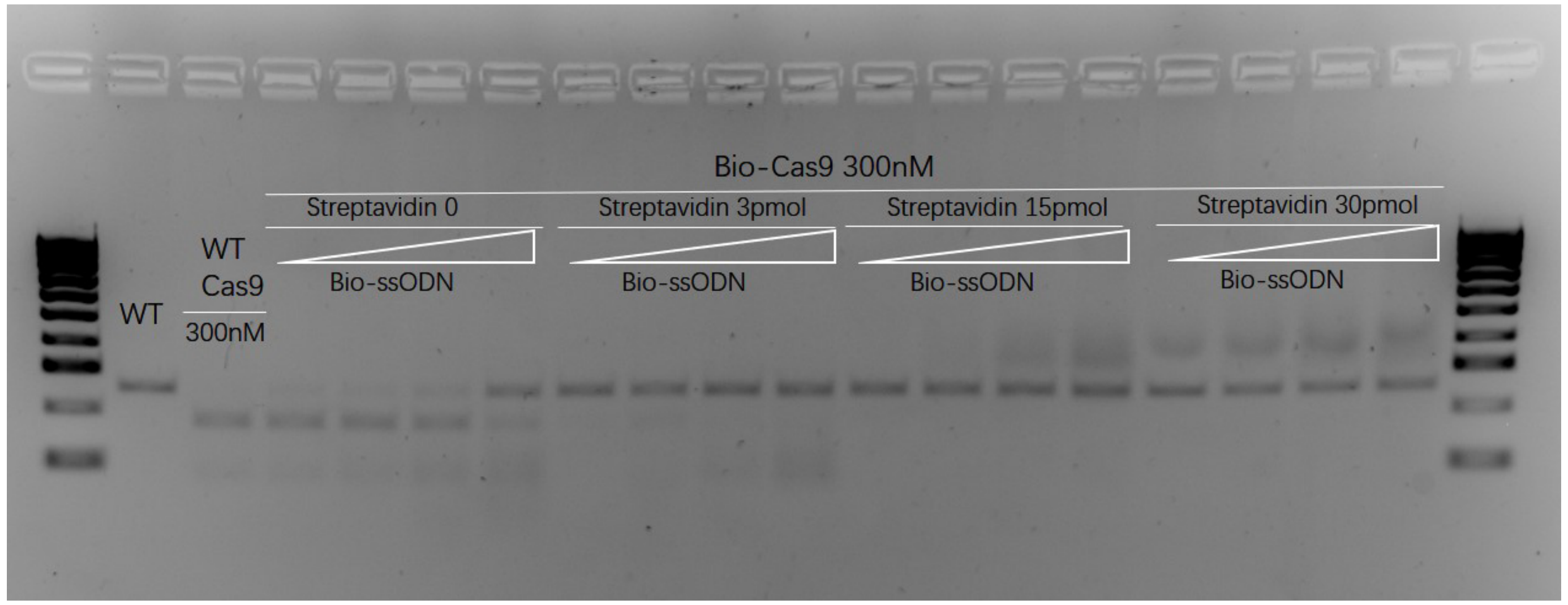


**B Cell lines monitored for mycoplasma after editing**

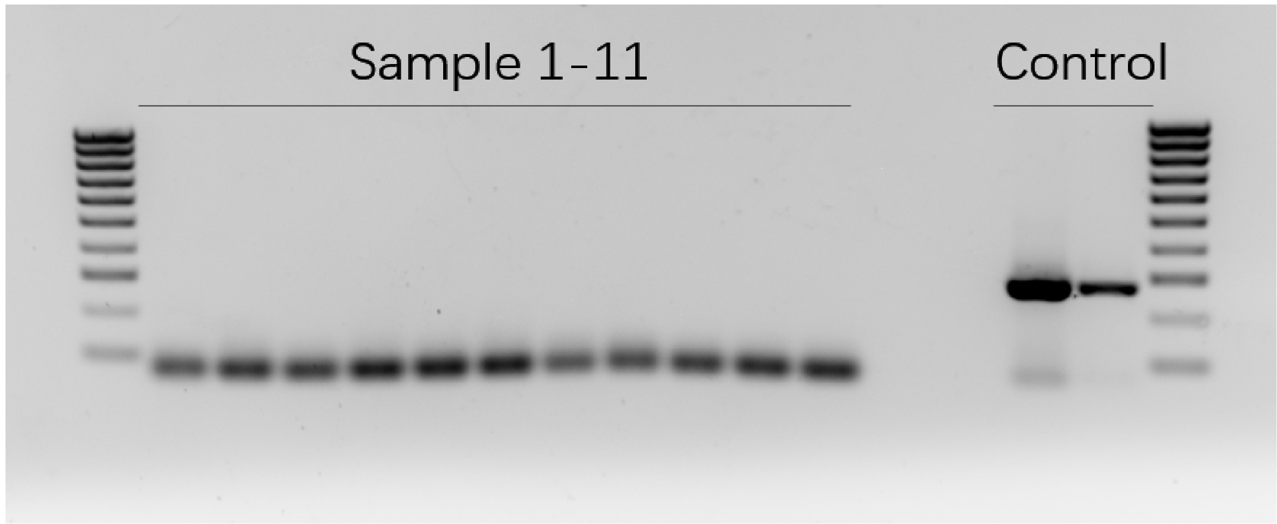


Supplmentary Figure S6
